# Single-cell response to Wnt activation in human embryonic stem cells reveals uncoupling of Wnt target gene expression

**DOI:** 10.1101/2023.01.11.523587

**Authors:** Simon Söderholm, Amaia Jauregi-Miguel, Pierfrancesco Pagella, Valeria Ghezzi, Gianluca Zambanini, Anna Nordin, Claudio Cantù

**Author notes:** These authors contributed equally to this work (Simon Söderholm, Amaia Jauregi-Miguel, Pierfrancesco Pagella, Gianluca Zambanini). Twitter: @ClaudioCantu81 (Claudio Cantù).

## Abstract

Wnt signaling drives nuclear translocation of β-catenin and its subsequent association with the DNA-bound TCF/LEF transcription factors, which dictate target gene specificity by recognizing Wnt responsive elements across the genome. β-catenin target genes are therefore thought to be collectively activated upon Wnt pathway stimulation. However, this appears in contrast with the non-overlapping patterns of Wnt target gene expression in several contexts, including early mammalian embryogenesis. Here we followed Wnt target gene expression in human embryonic stem cells after Wnt pathway stimulation at a single-cell resolution. Cells changed gene expression program over time consistent with three key developmental events: i) loss of pluripotency, ii) induction of Wnt target genes, and iii) mesoderm specification. Contrary to our expectation, not all cells displayed equal amplitude of Wnt target gene activation; rather, they distributed in a continuum from strong to weak responders when ranked based on the expression of the target *AXIN2*. Moreover, high *AXIN2* did not always correspond to elevated expression of other Wnt targets, which were activated in different proportions in individual cells. This uncoupling of Wnt target gene expression, which was also identified in single colorectal cancer cells with hyper-activated Wnt signaling, underlines the necessity to identify additional mechanisms that explain the heterogeneity of the Wnt/β-catenin-mediated transcriptional outputs in single cells.

## Introduction

Wnt/β-catenin signaling is an evolutionarily conserved mechanism for cell-cell communication that induces a change in the behavior of signal-receiving cell by activating a specific, Wnt-induced gene expression program (Valenta et al., 2012). In the current model, free cytoplasmic β-catenin is rapidly marked for degradation by a cytoplasmic destruction complex that consists of AXIN, the adenomatous polyposis coli (APC) tumour suppressor protein, glycogen synthase kinase 3 (GSK3α and GSK3β, both referred to as GSK3) and casein kinase 1 (CK1) (Gammons and Bienz, 2018; Lyou et al., 2017). In the signal receiving cells, WNT ligands or pharmacological blockage of GSK3 cause the inhibition of the destruction complex, leading to the accumulation and nuclear translocation of β-catenin (Li et al., 2012; MacDonald et al., 2009; Wu et al., 2013). In the nucleus, β-catenin binds to the T-cell factor/lymphoid enhancer factor (TCF/LEF) family of transcription factors, and activates the transcription of Wnt target genes (Cadigan and Waterman, 2012). TCF/LEF recognize a select consensus sequence on the DNA present within genomic <u>W</u>nt <u>r</u>esponsive <u>e</u>lements (WREs) and thereby impose specificity as to which genes are activated by the WNT/β-catenin biochemical cascade (Cadigan and Ramakrishnan, 2017). While the existence of tissue-specific Wnt-dependent transcriptional outcomes has been proposed (Afouda et al., 2020; Cantù et al., 2018; Mukherjee et al., 2020; Nakamura and Hoppler, 2017; Nakamura et al., 2016; Pagella et al., 2022; Ramakrishnan et al., 2022), to the best of our knowledge no investigation has been conducted on the single-cell specificity of the Wnt transcriptional response.

Common readouts of Wnt activation include mRNA measurements (such as the widely used target genes *AXIN2* and *SP5*, (Jho et al., 2002a; Weidinger et al., 2005)) or the use of transcriptional reporters (e.g., *TOPflash* in vitro or *BATgal* in vivo (Maretto et al., 2003)). In this study we inquired whether these assays have been measuring the average response of a cell population, while individual cells might possess diverse behaviors. Understanding this would reveal if the use of individual target genes or reporters constitute reliable “shortcuts” to assess that global activation of the Wnt-mediated transcriptional program has occurred. We addressed this question by following Wnt target gene activation in human embryonic stem cells (hESCs) via single cell RNA-sequencing (scRNAseq) after GSK3 inhibition. In these conditions, hESCs activate Wnt signaling, depart from a pluripotent state and undergo mesodermal differentiation, thereby recapitulating key steps of lineage commitment during human gastrulation (Davidson et al., 2012). We discovered that during this process cells vary in the extent to which they upregulate *AXIN2*, and that they can be ranked in a distribution ranging from strong to weak responders.

While the bulk population upregulated several target genes concomitantly, at the single cell level we noticed that high *AXIN2* mRNA did not necessarily correspond to elevated expression of other Wnt targets, such as *SP5*. Vice versa, several cells displaying high *SP5* did not upregulate *AXIN2*. This observation allowed us to distinguish different Wnt-responsive cell subpopulations, within an otherwise seemingly homogeneous cell population, based on their relative proportions of target gene activation. We refer to the existence of these subpopulations as “uncoupling” of Wnt target gene expression. This phenomenon suggests that individual cells must possess diverse mechanisms to activate cell-specific Wnt-dependent transcriptional programs. In light of this, we submit that assays based on individual reporter genes or constructs can at best monitor the average response to Wnt signaling and might fail to reveal alternative yet equally relevant gene expression programs. Moreover, our study underscores the need to discover how the TCF/LEF/β-catenin transcriptional complex generates heterogeneous transcriptional outcomes within single cells.

## Results

### Detection of Wnt-dependent transcriptional response at single-cell resolution

We wondered whether monitoring individual target genes or transcriptional reporters is a reliable method to assess the response to Wnt signaling or, alternatively, if this measurement constitutes the average of a spectrum of cell-specific responses. To discern between these two possibilities, we stimulated human embryonic stem cells (hESCs) with the Wnt signaling pharmacological agonist CHIR, which inhibits the negative pathway regulator GSK3 (Wu et al., 2013). As previously reported, continuous stimulation of hESCs with CHIR activates Wnt/β-catenin signaling, drives exit from pluripotency, and the acquisition of mesodermal identity (Davidson et al., 2012). At defined time-points (Pagella et al., 2022) we assessed the dynamics of the transcriptional program via single-cell RNA sequencing (scRNAseq) (Figure 1A). We envisioned two possible outcomes: either cells would respond homogeneously, namely each cell would upregulate Wnt target genes to the same extent, or heterogeneously, if our measurement identified different levels of target gene expression across individual cells (Figure 1B). We quantified the mRNA content of a total of 12,798 hESCs, distributed over four time points: 0 hours (h; this is our Wnt-OFF baseline); 4h; 24h; 72h (Figure S1A). Visualization by uniform manifold approximation and projection (UMAP) (McInnes et al., 2018) shows that, when stimulated, hESCs globally change identity starting from 24h (Figure 1C). Global gene expression analysis confirmed a broad shift in the transcriptomic signature of these populations, characterized by the loss of expression of pluripotency markers and the gradual appearance of differentiation and mesodermal traits (Figure 1D). We extrapolated genes that are relevant for these developmental processes and ranked them according to their known role (Figure 1E). For instance, *POU5F1*/*OCT4, SOX2* and *MYC* are broadly (that is, in a large fraction of the population) and highly (presenting high read counts per cell) expressed in hESCs at 0h, and both these parameters progressively decrease (Figure 1E, upper panel). Concomitantly and as expected, differentiation markers such as *TBXT* (also known as *T*/*Brachyury*) and *MIXL1*, which are not present at 0h, only appear after differentiation have been induced by GSK3 inhibition (Figure 1E, lower panel). These observations were in line with current knowledge (Davidson et al., 2012; Efroni et al., 2008; Tajonar et al., 2013), and constituted an important validation of our new dataset.

**Figure 1.**
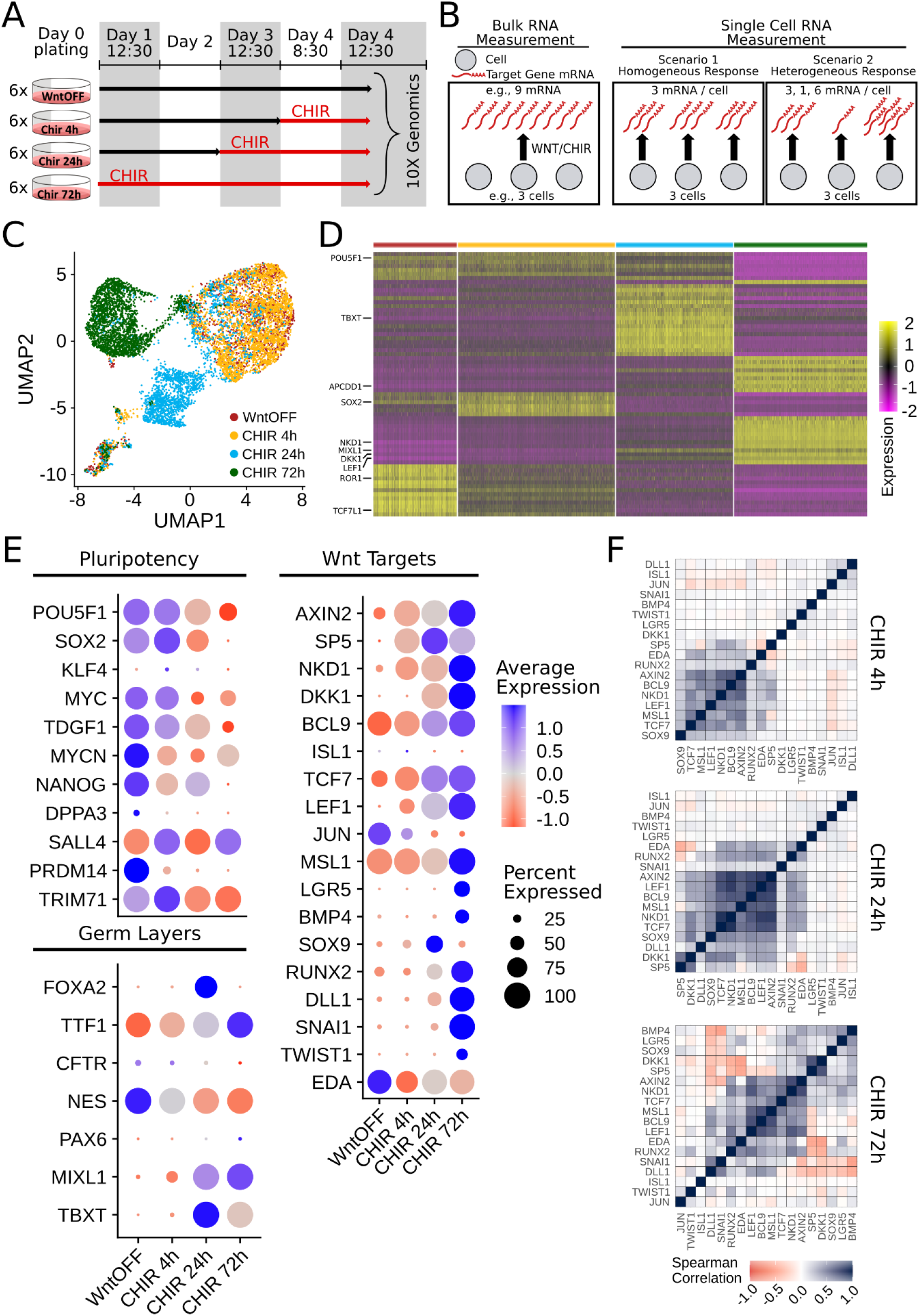
Time-resolved analysis of Wnt-dependent transcriptional response at single-cell resolution. (A) Experimental approach. Wnt/β-catenin signaling was activated in human embryonic stem cells (hESCs) by treatment with the GSK3B inhibitor CHIR99021 (CHIR). hESCs were collected at 4 hours (4h), 24 hours (24h) and 72 hours (72h) after Wnt activation and processed for single-cell RNA sequencing (10X genomics pipeline). For each time point, six independent wells were pooled and sequenced. (B) Wnt stimulation induces the expression of Wnt target genes at a bulk population level. However, this global phenomenon could be the consequence of two very different single-cell scenarios: i) cells respond homogeneously, namely each cell would upregulate Wnt target genes to the same extent, or ii) heterogeneously, i.e., different cells activate diverse sets of target genes. (C) UMAP visualization of the four datasets (WNT-OFF, 4h, 24h, 72h) merged and harmonized. The color code indicates the different time points. (D) Heatmap showing scaled normalized expression of genes differentially expressed between the time points analyzed. For the heatmap visualization the top 25 genes differentially expressed for each time point compared to all other time point were selected. (E) Dot-plot showing the expression of pluripotency markers, differentiation markers, and Wnt target genes over the time of Wnt/β-catenin activation. Color scale indicates the relative mRNA expression level, while the dot diameter is proportional to the fraction of cells that express each gene. (F) Spearman correlation plots showing the single-cell-level correlation between the expression of each Wnt target gene at the different time points.

We were curious to track the time-dependent expression of Wnt target genes. We thus selected a curated list of several historically characterized Wnt targets (https://web.stanford.edu/group/nusselab/cgi-bin/wnt/target_genes) (Figure 1E, right panel). Some of these increased their expression upon CHIR treatment (*BCL9, MSL1, AXIN2*), while others only started to be expressed at 4h (*SP5, LEF1, NKD1*) or at later time points (*DKK1, RUNX2, SNAI1*). We considered this behavior notable as it suggested the existence of early, intermediate, and late phases of Wnt pathway activation. We calculated the Spearman Correlation between the expression of all these targets, which allowed us to pinpoint that a subset of these genes positively correlated at each time point of analysis, in line with current knowledge (blue squares in Figure 1F). Interestingly, a significant fraction of targets showed mild, absent, or even inverse correlation (light blue, white, red squares in Figure 1F, respectively). This would indicate that all target genes were not necessarily concomitantly activated upon Wnt pathway stimulation, and thus that mechanisms to uncouple their regulation must exist: we refer to this phenomenon as “uncoupling” of Wnt target gene expression. We submit that the current understanding of how Wnt target genes are mechanistically transcribed fails to explain this uncoupling observed at single cell resolution.

### Uncoupling of Wnt target gene expression in single cells

UMAP projection allowed us to visualize the expression intensity of each target gene in all cells. Current knowledge would imply that either a cell would present activated Wnt pathway (Wnt-ON), detectable by increased expression of all targets concomitantly, or it would not (Wnt-OFF). While UMAPs showed cells respecting this pattern, it also revealed that target gene expression was not homogeneously distributed across the cell population (Figure 2A). While the four prototypical targets *AXIN2, SP5, NKD1* and *LEF1* displayed a considerable overlap in their expression pattern, there were also cell populations characterized by different signal intensities for these genes (Figure 2A). Our dataset allowed us to explore this phenomenon by ranking the cells based on their relative expression of a specific gene, used as a reporter to monitor the Wnt activation status in individual cells. We selected *AXIN2*, which is frequently used as representative proxy of the Wnt/β-catenin transcriptional response (Jho et al., 2002b) (Figure 2B). *AXIN2* was almost undetectable or expressed at very low levels in Wnt-OFF, and its expression progressively increases at all subsequent time points (Figure 2B). However, contrary to the simplistic prediction that all cells should equally upregulate *AXIN2* upon CHIR treatment, individual hESCs displayed highly variable read-counts mapped on the *AXIN2* coding sequence: single hESCs were found along a continuous curve, a distribution than underlies the existence of strong and weak responders to Wnt pathway stimulation. This *AXIN2* gradual slope is most likely not a consequence of the sparsity of reads that is often observed in scRNAseq experiments (Pollen et al., 2014); this is demonstrated by the remarkable overlap between *AXIN2* and *NKD1* curves which hinted that strong responders similarly upregulated both these genes, while weak responders do it to a lesser extent (Figure 2B).

**Figure 2.**
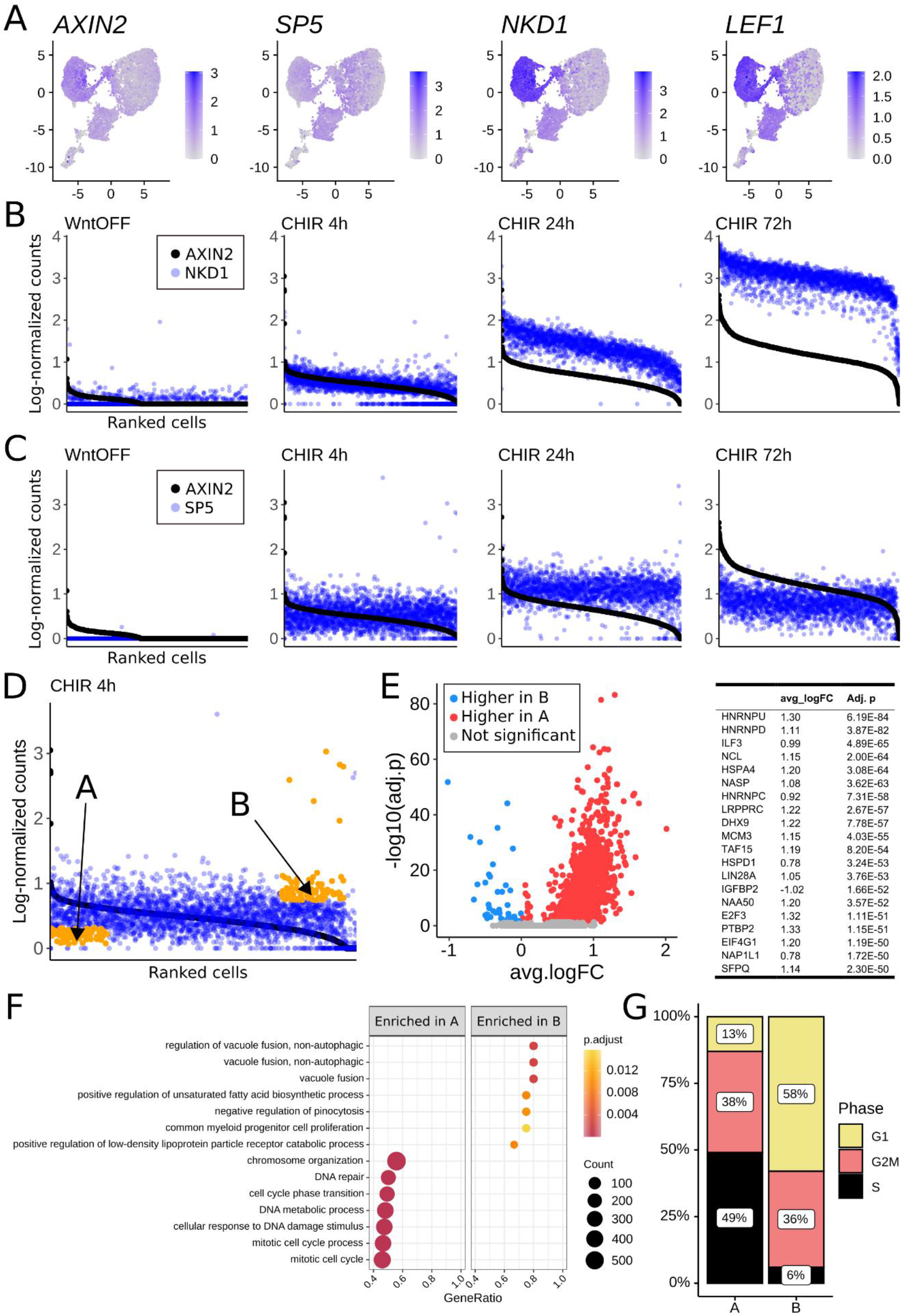
Uncoupling of Wnt target gene expression at the single-cell level. (A) Feature plots showing the expression of the Wnt target genes *AXIN2, SP5, NKD1* and *LEF1* in hESCs in WNT-OFF conditions and after 4 hours, 24 hours, and 72 hours of Wnt / β-catenin signaling activation (UMAP plot color-coded for time-points: Figure 1C). Notice that expression of Wnt target genes did not completely overlap. (B) Single-cell *NKD1* mRNA expression (blue dots) represented by ranking cells based on their *AXIN2* mRNA expression (black dots; cells distributed on the horizontal axis, from *AXIN2*-highest on the left to *AXIN2*-lowest on the right). *NKD1* and *AXIN2* expression were highly correlated: both *AXIN2* and *NKD1* expression increased over time of Wnt stimulation, and high *AXIN2* expression was accompanied by high *NKD1* expression at a single cell level at any time point. (C) Single-cell *SP5* mRNA expression (blue dots) represented by ranking cells based on their *AXIN2* mRNA expression (black dots, ordered as in B). *SP5* and *AXIN2* expression were uncoupled on a temporal basis: while both *SP5* and *AXIN2* were upregulated in response to Wnt stimulation until 24 hours, *SP5* was downregulated after 72 hours, while *AXIN2* expression continued to increase. (D) *AXIN2* and *SP5* expression was uncoupled among single cells stimulated for 4 hours. Already at this early time point, it was possible to identify distinct populations of *AXIN2*^*high*^ */ SP5*^*low*^ and *AXIN2*^*low*^ */ SP5*^*high*^ hESCs (orange dots - A and B, respectively) Cells in A were selected by extracting the top 20% *AXIN2*-expressing cells and from these select 100 cells with the lowest expression of *SP5*. Likewise, for B, the bottom 20% of *AXIN2*-expressing cells were extracted and from these 100 cells with the highest expression of *SP5* were selected. (E) Volcano plot and table showing differentially expressed genes (Fc > 0.25; adj. p-value < 0.05) between *AXIN2*^*high*^ */ SP5*^*low*^ (“A” in panel D) and *AXIN2*^*low*^ */ SP5*^*high*^ (“B” in panel D). Red dots: genes expressed at significantly higher levels in A; blue dots: genes expressed at significantly higher levels in B. The table displays the top 20 significantly differentially expressed genes. (F) Gene set enrichment analysis of genes differentially expressed between *AXIN2*^*high*^ */ SP5*^*low*^-A and *AXIN2*^*low*^ */ SP5*^*high*^-B hESCs. *AXIN2*^*high*^ */ SP5*^*low*^-A showed higher expression of genes involved in the cell cycle progression, while *AXIN2*^*low*^ */ SP5*^*high*^-B displayed a transcriptomic signature typical of metabolically active cells. (G) *AXIN2*^*high*^ */ SP5*^*low*^-A and *AXIN2*^*low*^ */ SP5*^*high*^-B populations showed a different distribution in the cell cycle. While *AXIN2*^*high*^ */ SP5*^*low*^-A cells were mostly in G2/M and S-phase, in the *AXIN2*^*low*^ */ SP5*^*high*^-B subpopulation the number of cells in G1 phase grew considerably at the expense of cells in S phase.

We overlaid the expression intensity per cell of another notable Wnt target gene, *SP5* (Huggins et al., 2017), on the *AXIN2*-expression-based distribution. *SP5* was not expressed in Wnt-OFF, but consistent with it being a Wnt/β-catenin target (Huggins et al., 2017), it was promptly and broadly transcribed at 4h (Figure 2C). From the comparison between *AXIN2* and *SP5* expression, two important observations emerged. First, as previously hinted, there was a temporal disconnect in the transcription of the two target genes: while *AXIN2* levels continued to rise until 72h, *SP5* peaked at 24h and decreased at the later time point. Second, the distribution of *SP5* expression in single cells did not closely follow that of *AXIN2*. Several cells that could be defined as strong responders due to high *AXIN2* showed little to no levels of *SP5* (*AXIN2*^*high*^; *SP5*^*low*^); conversely, numerous presumed weak responders (low in *AXIN2*) displayed high expression of *SP5* (*AXIN2*^*low*^; *SP5*^*high*^) (Figure 2C). This posits a challenge in defining what cell populations are Wnt signaling responsive: the conclusion would change if the experimenters were to choose *AXIN2* or *SP5*. We submit that this conundrum demands primary attention from the field studying ontogenetically relevant signaling pathways. Notably, this phenomenon is not restricted to *AXIN2* and *SP5*; a similar observation was made concerning the relationship between *AXIN2* and several other target genes (Figure S2).

To test if the uncoupling of *AXIN2* and *SP5* reflected a real difference in cell populations, we informatically isolated *AXIN2*^*high*^; *SP5*^*low*^ and *AXIN2*^*low*^; *SP5*^*high*^ hESC subpopulations at the earliest time point (4h; Figure 2D) and performed differential gene expression analysis (Figure 2E). The selection of the earliest time point permitted us to focus on direct transcriptional effects occurring soon after GSK3 inhibition, thereby diminishing the relative contribution of secondary effects downstream of GSK3 activity, which include feedback loops and Wnt-STOP (Acebron et al., 2014; Koch et al., 2015; Taelman et al., 2010). Our analysis revealed that these two populations are classified across different stages of the cell cycle. In particular, the *AXIN2*^*low*^; *SP5*^*high*^ sub-population showed a high fraction of cells in S-phase, indicative of cell proliferation (Figure 2F, G; population A in figure 2D). The *AXIN2*^*high*^; *SP5*^*low*^ sub-population instead displayed a largest fraction in G1 phase, typical of metabolically active cells (Figure 2F, G; population B in Figure 2D). This underscores a physiological difference between the two *in silico* selected cell populations and supports the notion that the uncoupling of Wnt target genes is associated with existing cellular heterogeneity. One limitation of our analysis is that it did not allow us to distinguish whether a different responsiveness to Wnt drives changes in the cell cycle pattern, or if cells at different cell cycle stages exhibit differential response to Wnt. Future experiments are required to assess these possibilities. Nonetheless, our finding was strikingly reminiscent of a phenomenon previously referred to as mitotic Wnt signaling (Davidson et al., 2009). We speculate that the *AXIN2*^*high*^; *SP5*^*low*^ population – several of which are in G1 – might predominantly include cells which had a mitotic peak of Wnt signaling, and that are subsequently found in the successive phase of the cell cycle.

### Physical β-catenin targets are differentially regulated in single cells

The previous analysis indicated that Wnt signaling drives the selective activation of subsets of targets in different cells. To test if the uncoupling is a consequence of direct regulation by β-catenin, we integrated the scRNAseq analysis with the genome-wide physical occupancy of β-catenin obtained via Cut Under Targets & Release Upon Nuclease Low Volume Urea (CUT&RUN-LoV-U; (Zambanini et al., 2022)) at matching time points (Pagella et al., 2022). This exposed groups of direct physical Wnt/β-catenin genomic targets based on the timing of β-catenin association with their locus. We simultaneously plotted their relative expression (color coded, Figure S3) and the fraction of cells expressing each specific target (dot size, Figure S3). Among all targets, several display high correlation between timing of expression and of β-catenin binding, emphasizing the causal involvement of β-catenin association to the chromatin in their transcriptional regulation (Figure 3A). For example, early physical targets (e.g., *AXIN2, SP5*) are promptly upregulated, albeit, according to our previous observation (Figure 2), in heterogeneous amounts across single cells (Figure 3B). By the same token, late physical targets (e.g., *DKK1, ZNF703*) are transcribed at 72h (Figure 3A, C). The integration of these two technologies provides a reliable system to pinpoint the time-dependent correlation in the expression of direct Wnt/β-catenin physical targets.

**Figure 3.**
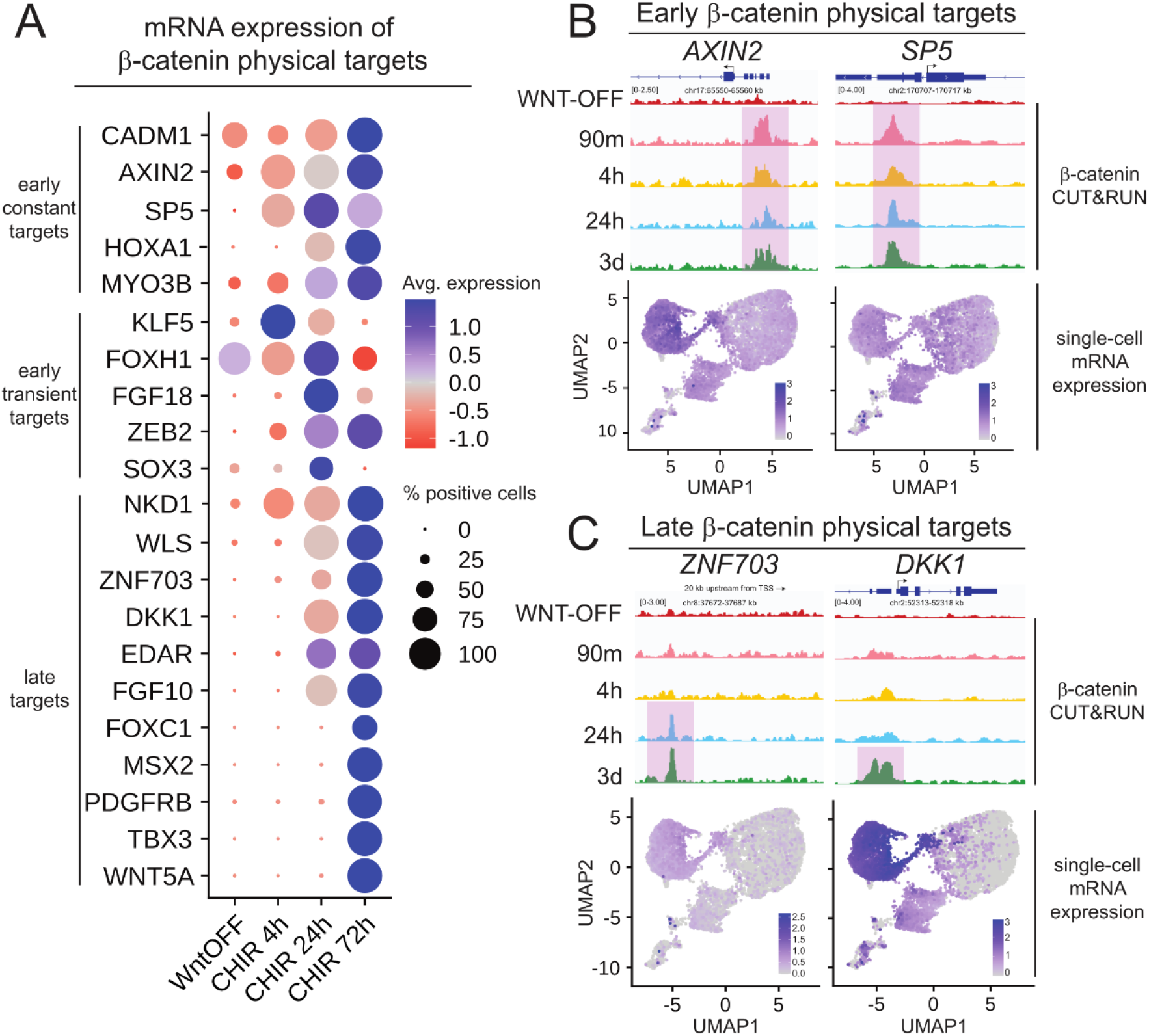
Correlation between bulk β-catenin physical genomic targets and mRNA expression at the single-cell level. (A) Dot-plot showing mRNA expression of genes whose loci were physically bound by β-catenin, as detected by CUT&RUN-LoV-U (Pagella et al., 2022). “Early constant targets” indicates loci bound by β-catenin already after 90 minutes of Wnt signaling activation and that remain targeted at all time points analyzed until 72 hours. “Early transient targets” indicates loci transiently bound by β-catenin after 90 minutes or 4 hours of stimulation. “Late targets” are loci targeted by β-catenin only after 24 hours of stimulation or more. Color scale indicates the relative mRNA expression level, while the dot diameter is proportional to the fraction of cells that express each gene. (B) Correlation between the β-catenin genomic binding profile (upper row; β-catenin binding is highlighted by pink box) at the early, constant target loci *AXIN2* and *SP5*, and their mRNA expression at single cell level (feature plots, lower row). Both genes were expressed throughout the time points analyzed, but their relative distribution diverged significantly. (C) Correlation between the β-catenin genomic binding profile (upper row; β-catenin binding is highlighted by pink box) at the late target loci *ZNF703* and *DKK1*, and their mRNA expression at single cell level (feature plots, lower row). *ZNF703* and *DKK1* expression was concentrated in cells stimulated for 72 hours, in accordance with the late recruitment of β-catenin.

### Wnt target genes are uncoupled in Wnt-driven human colorectal cancers

We were not satisfied in observing such a surprising phenomenon – the uncoupling – in only one experimental set-up. Therefore, we set out to investigate whether Wnt target genes are uncoupled in other physiologically relevant contexts. We interrogated the expression of Wnt target genes across single cells from healthy human intestinal epithelium and tumor deriving from this tissue (that is, colorectal cancer, CRC; (Uhlitz et al., 2021)). For both tissues, we focused on epithelial cells (*VIL1*-positive) and on the subset of crypt-residing intestinal stem cells (*LGR5*-positive) (Figure 4A). The latter in particular is thought to be a homogeneous population exquisitely dependent on Wnt signaling (Sato et al., 2011). Spearman correlation of Wnt target gene expression on a population level revealed, as shown above for hESCs, instances of positive correlation (blue-graded signal in Figure 4B) as well as uncoupling, which includes both absence of and weak (positive/negative) correlation (light blue-white-light red-graded signal in Figure 4B). CRC cells showed higher correlation within a set of reliable Wnt targets (Figure 4C); we consider this consistent with a general constitutive activation of Wnt signaling in CRC (van Kappel and Maurice, 2017; Lyou et al., 2017), and an important internal validation of our analysis. The uncoupling is also observed at a single-cell resolution: both *VIL1*-positive and *LGR5*-positive cells did not display matching distributions of *AXIN2* and *SP5* mRNA (Figure 4C). This is particularly remarkable when considering the LGR5-positive cells from the CRC subset, which likely represent the cancer stem cells whose renewal is dependent on constitutively active Wnt/β-catenin pathway (Groden et al., 1991; Kinzler et al., 1991).

**Figure 4.**
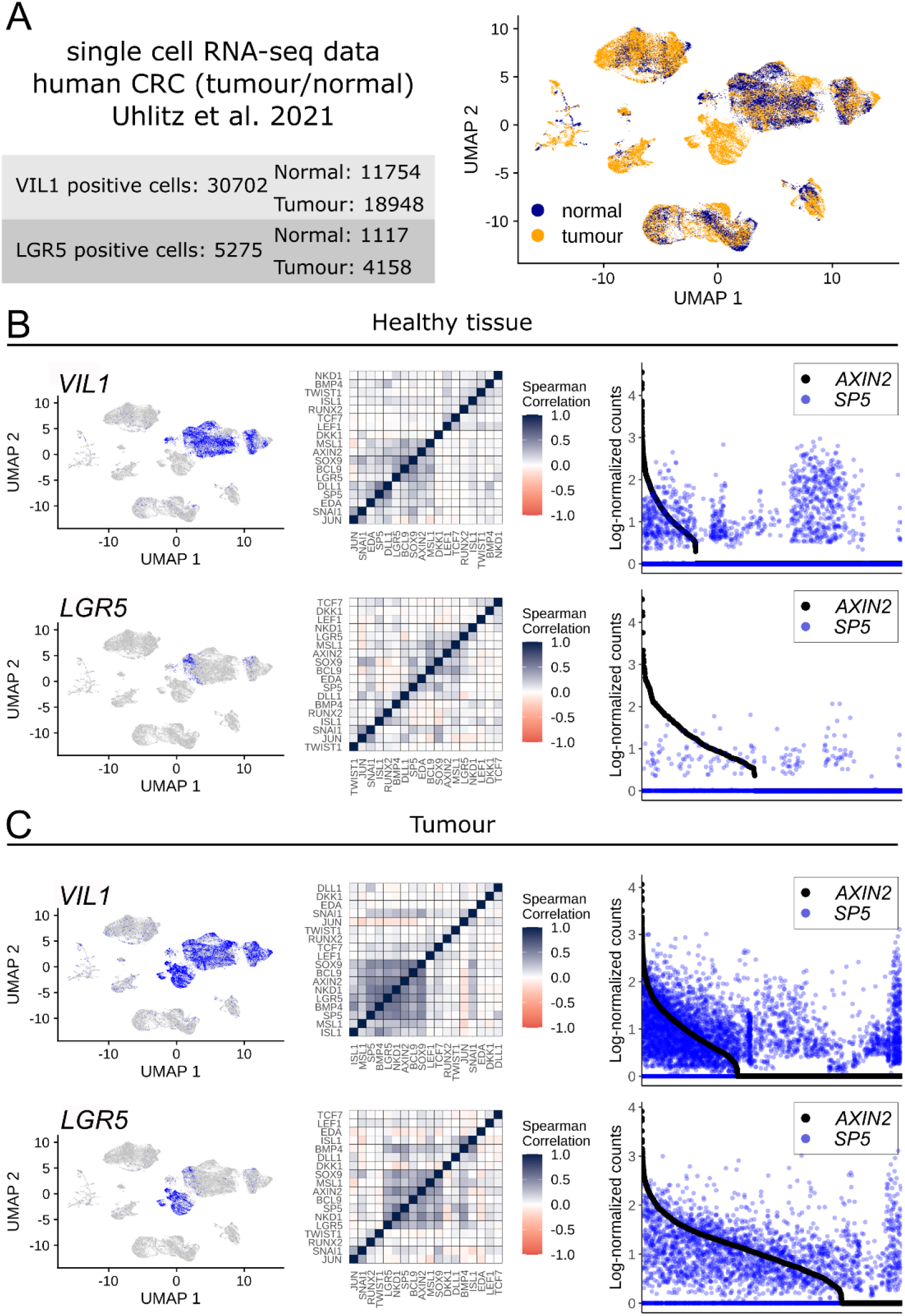
Uncoupling of Wnt target gene expression in human colorectal cancer (CRC). (A) Overview and UMAP representation of the single-cell atlas of human colon and human colorectal cancer, from (Uhlitz et al., 2021). Colors in the UMAP distinguish cells from the healthy colon (blue) and from CRC (yellow). (B) Correlation and uncoupling of Wnt target gene expression in cells from healthy colon epithelium. Left column: feature plots showing the distribution of *VIL1* (intestinal epithelial cells marker) and *LGR5* expression (intestinal epithelial stem cells marker) in the dataset. Central column: heatmaps of Spearman correlation values between the expression of Wnt target genes. Right column: single-cell *SP5* mRNA expression (blue dots) represented by ranking cells based on their *AXIN2* mRNA expression (black dots; cells distributed on the horizontal axis, from *AXIN2*-highest on the left to *AXIN2*-lowest on the right). Wnt target gene expression showed limited correlation (Spearman correlation heatmaps, central column) within all colon epithelial cells (*VIL1*-positive; upper row) and within the *LGR5*-positive stem cells subset (lower row). At the single-cell level, we could detect numerous cells that showed uncoupled *AXIN2* and *SP5* expression (right column). (C) Correlation and uncoupling of Wnt target gene expression in CRC epithelial cells. CRC epithelial cells, and their *LGR5*-positive subset both showed higher correlation (Spearman correlation heatmaps, central column) in the expression of several Wnt target genes when compared to their counterparts within the healthy tissue. A higher proportion of tumor cells (either *VIL1*-positive or *LGR5*-positive) expressed detectable levels of *AXIN2* when compared to healthy cells; at the single-cell level, numerous cells showed strong uncoupling of *AXIN2* and *SP5* expression (right column).

## Discussion

Here we traced the transcriptional response to Wnt pathway activation in hESCs at a single-cell resolution, while they undergo mesodermal differentiation. We observed that Wnt target genes induction is not a binary event – that is, responder cells either activate all genes or none. In contrast, we measured a continuous spectrum of behaviors ranging from strong to weak Wnt responses. Moreover, cells that could be defined as strong responders based on one gene (e.g., *AXIN2*) were not necessarily strong responders based on other direct targets (e.g., *SP5*): we define this previously unappreciated phenomenon *uncoupling* of Wnt target genes.

From a methodological and experimental perspective, this observation poses the problem of how to define Wnt responsive cells only based on individual reporters or on one/few genes (van Amerongen et al., 2012; Jho et al., 2002b; Maretto et al., 2003). In both datasets presented here, the search for strong Wnt responding individual cells would yield diverging results if based on, for example, one of the two targets *AXIN2* or *SP5*. This was unexpected, as these genes are both rapidly upregulated by Wnt/CHIR as their transcription is directly controlled by the TCF/LEF-β-catenin transcriptional complex (Huggins et al., 2017; Jho et al., 2002b). While the current model of Wnt-dependent transcription cannot account for these differences (van Tienen et al., 2017), sparse experimental evidence previously suggested that β-catenin might regulate alternative targets via diverse mechanisms, for example by recruiting cohorts of co-factors in a promoter-specific fashion (Brannon et al., 1997; Hecht et al., 2000). It is therefore imperative for the field to focus on the discovery of how individual Wnt target genes are mechanistically regulated. Moreover, we suggest caution when selecting the necessary reporter genes as “shortcut” for reporting Wnt activation.

While the uncoupling might appear as a surprise, it is also true that, when they are studied in depth, the regulatory mechanisms for the expression of individual genes can be convoluted, require the interplay of a complex syntax of regulatory regions (Bahr et al., 2018; Hörnblad et al., 2021) and integration from multiple signaling mechanisms (Hnisz et al., 2013). By the same token, also *AXIN2, SP5* as well as other genes typically classified as Wnt targets might be under the control of similar processes and their expression modulated by additional simultaneous drivers. This leads to the prediction that the proteins encoded by these genes might have novel, unknown functions in addition to being Wnt downstream effectors or regulators.

One key question remains to be answered, namely whether an initial heterogeneity present in a cell population deterministically influences the relative response to the same trigger, or the heterogeneity is generated as a consequence of the stimulus in a stochastic or statistically predictable manner. At this stage, we favor the first possibility where the constitution of individual cells matters for how they will respond. Indeed, the uncoupling of Wnt target gene expression was observed after short intervals of time, largely before the establishment of feedback loops and other secondary consequences in cascade. Therefore, we propose that hESCs and other Wnt responsive cells that display the uncoupling must possess an intrinsic heterogeneity from which a spectrum of responses is established. Consistent with this view, the Pelkmans group recently discovered that single cells responded heterogeneously to equal concentrations of EGF likely by combining cellular-state and context-dependent stimuli (Kramer et al., 2022). As EGF signaling, also Wnt and other developmentally relevant signaling pathways might have evolved to include context-awareness when fostering cellular decision-making avenues.

## Acknowledgments

This work was supported by Grants to C.C. from Cancerfonden (CAN 2018/542 and 21 1572 Pj), the Swedish Research Council, Vetenskapsrådet (2021-03075), and Linköping University. C.C. is a Wallenberg Molecular Medicine (WCMM) fellow and receives generous financial support from the Knut and Alice Wallenberg Foundation. P.P. is supported by a fellowship awarded by the WCMM at Linköping University.

## Authors contributions

**Simon Söderholm:** Conceptualization, Methodology, Investigation, Formal Analysis, Data Curation, Writing – Review & Editing, Visualization. **Amaia Jauregi-Miguel:** Conceptualization, Methodology, Investigation, Writing – Review & Editing. **Pierfrancesco Pagella:** Conceptualization, Methodology, Investigation, Writing – Original Draft, Writing – Review & Editing. **Valeria Ghezzi:** Writing – Review & Editing. **Gianluca Zambanini:** Writing – Review & Editing. **Anna Nordin:** Writing – Review & Editing. **Claudio Cantù:** Conceptualization, Writing – Original Draft, Writing – Review & Editing, Supervision, Project Administration, Funding Acquisition.

## Supplementary Figures

**Figure S1.**
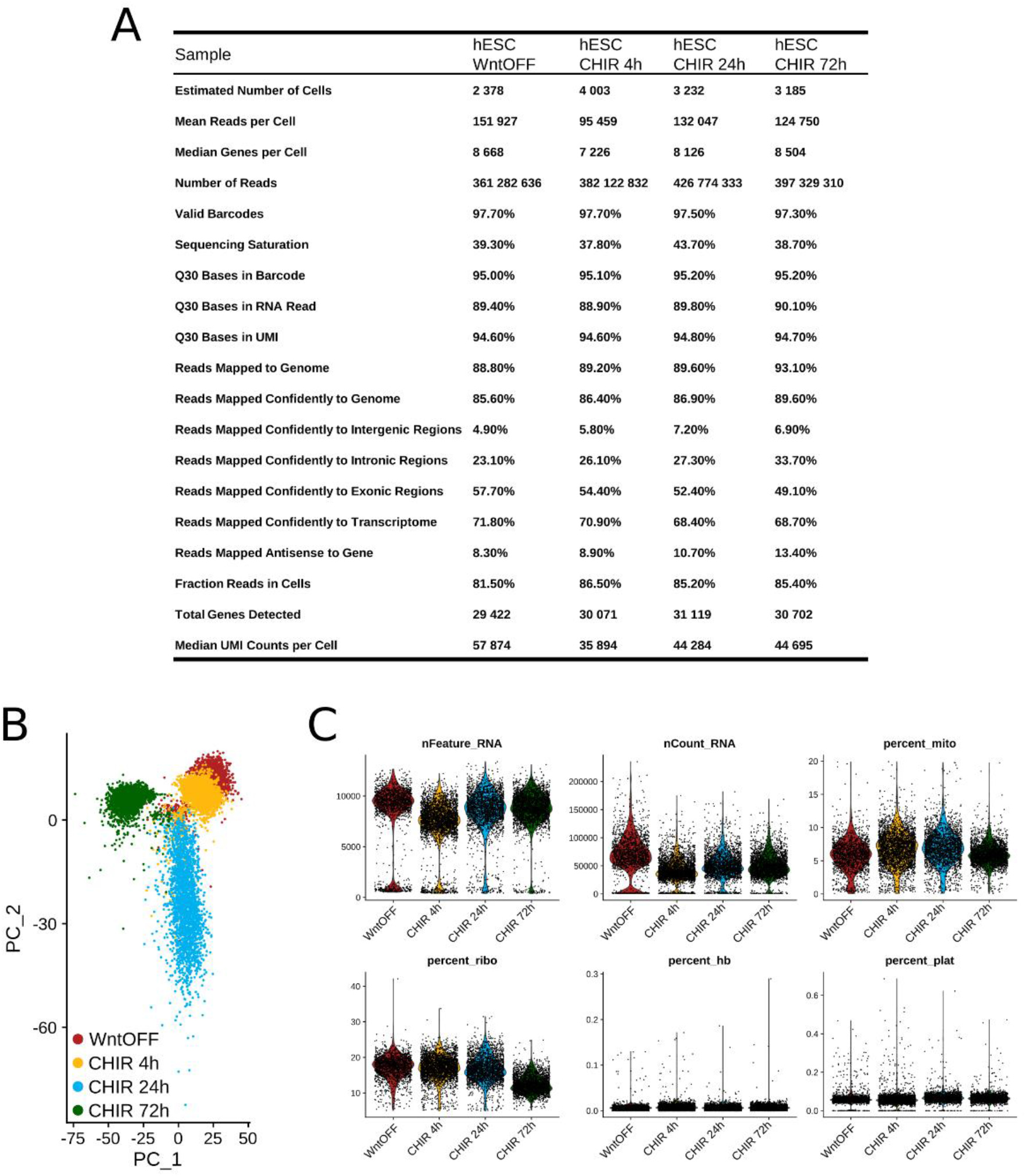
Quantitative details and quality control of single-cell data. (A) Samples size and sequencing outcome details. (B) Principal component analysis (PCA) groups samples based on treatment time. (C) Violin plots illustrate distribution of percentage of mitochondrial genes (mito), number of UMI counts (nCount) and number of genes with at least one UMI count (nFeature) per cell after subsetting cells according to the following quality control measures: cells with a percentage of mitochondrial read counts above 20% and/or below 5% ribosomal read counts were excluded, as well as cells with less than 200 genes.

**Figure S2.**
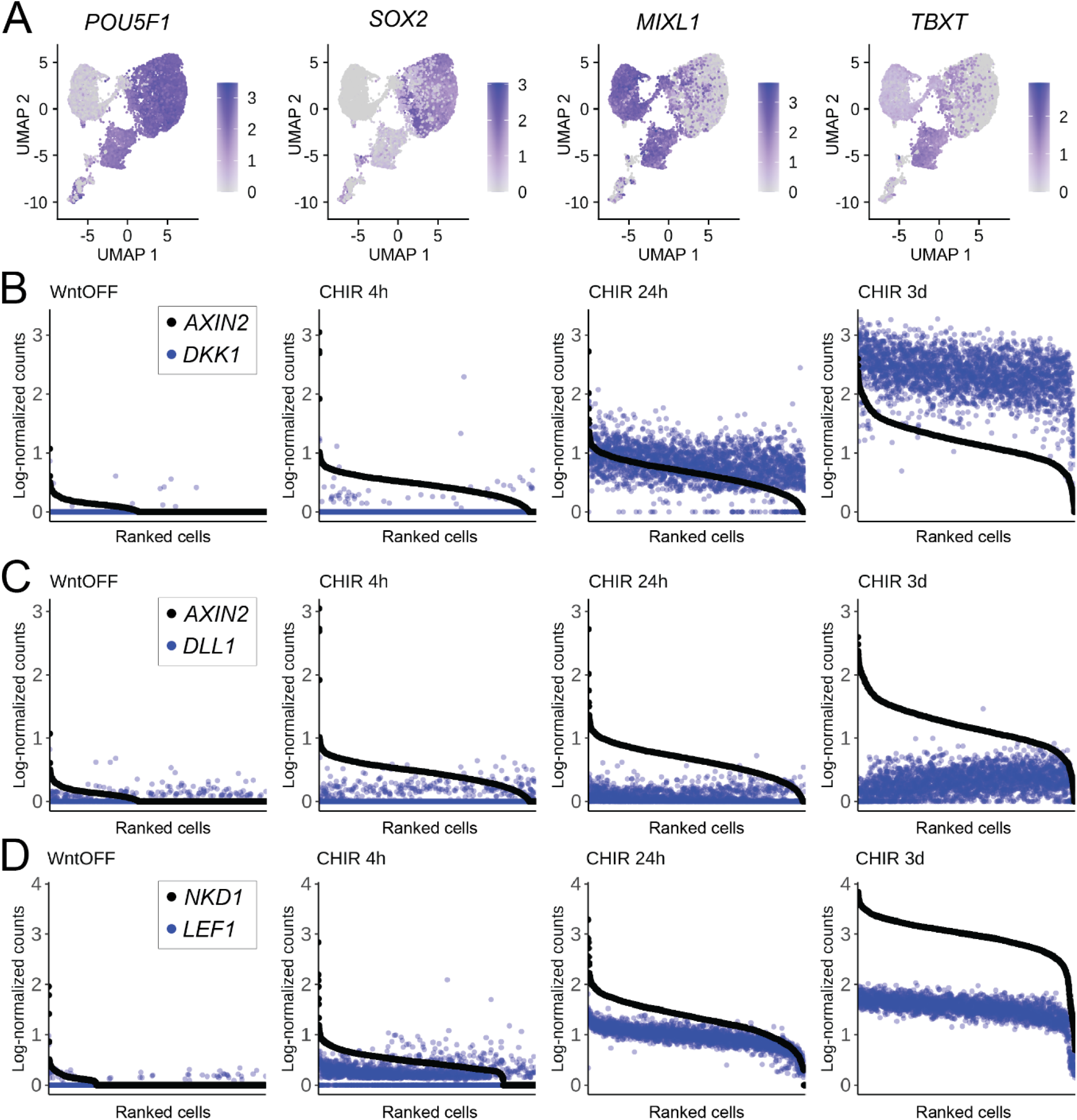
(A) Feature plots showing the expression of the pluripotency markers *POU5F1* and *SOX2*, and the mesoderm markers *TBXT* (early mesoderm) and *MIXL1* (early and late mesoderm) in hESCs. Expression of pluripotency factors is high in WNT-OFF conditions and after 4 hours of Wnt activation, and progressively decreases to then disappear at 72 hours. Conversely, mesoderm markers expression gradually increases over time. (B) Single-cell *DKK1* mRNA expression (blue dots) represented by ranking cells based on their *AXIN2* mRNA expression (black dots; cells distributed on the horizontal axis, from *AXIN2*-highest on the left to *AXIN2*-lowest on the right). *DKK1* and *AXIN2* expression were partially correlated. Both *AXIN2* and *DKK1* expression increased over time of Wnt stimulation, but *AXIN2* expression was activated already at 4h, while *DKK1* started to appear at later time points. At 24h, clear uncoupling of *AXIN2* and *DKK1* expression within single cells could be observed. (C) Single-cell *DLL1* mRNA expression (blue dots) represented by ranking cells based on their *AXIN2* mRNA expression (black dots; cells distributed on the horizontal axis, from *AXIN2*-highest on the left to *AXIN2*-lowest on the right). Expression of *AXIN2* and *DLL1* was mostly uncoupled. (D) Single-cell *LEF1* mRNA expression (blue dots) represented by ranking cells based on their *NKD1* mRNA expression (black dots; cells distributed on the horizontal axis, from *NKD1*-highest on the left to *NKD1*-lowest on the right). Expression of *NKD1* and *LEF1* was highly correlated: high expression of *NKD1* was accompanied by high expression of *LEF1*.

**Figure S3.**
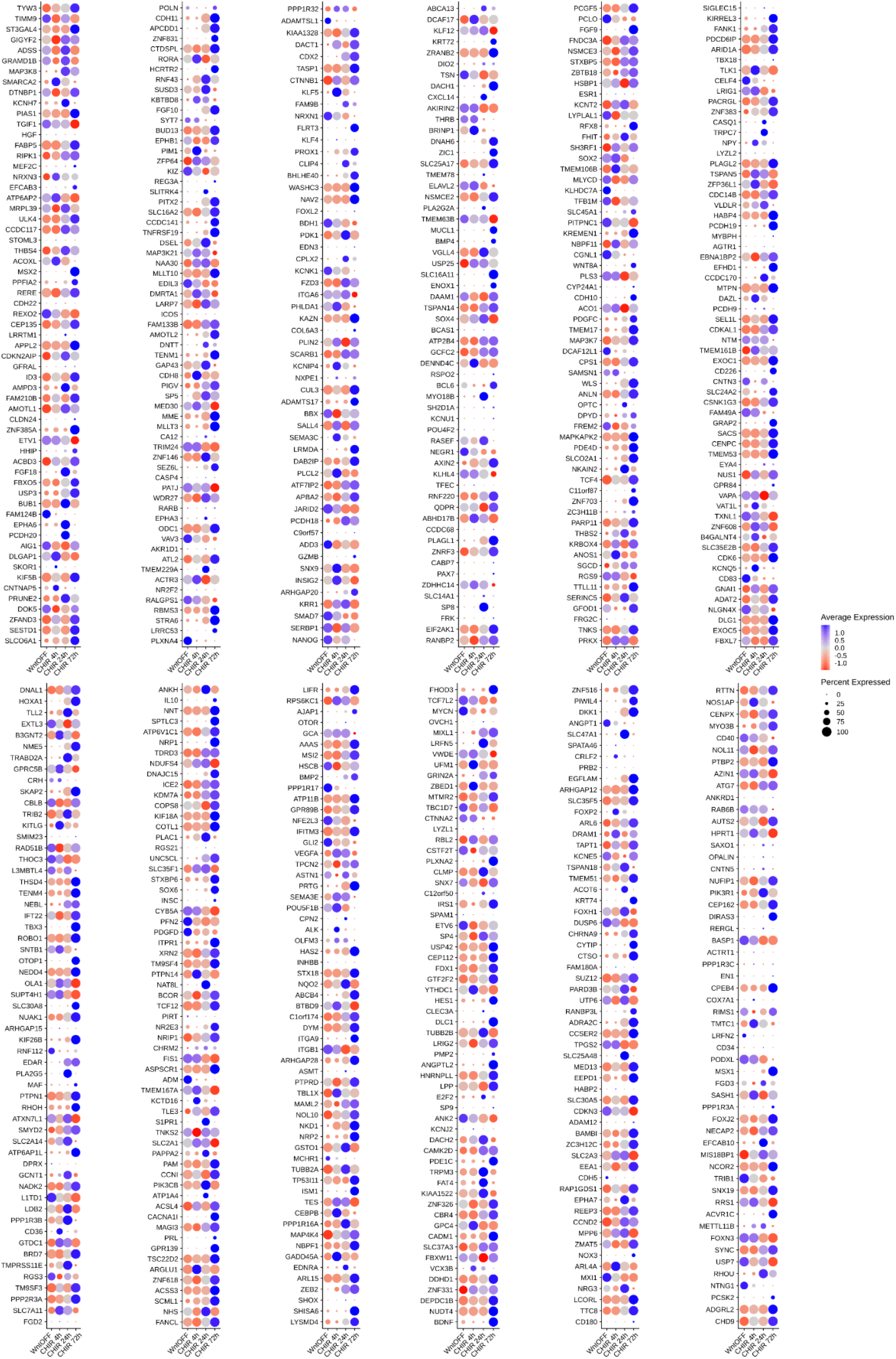
Time-resolved single-cell mRNA expression of β-catenin physical targets. Dot-plot showing mRNA expression of genes whose loci were physically bound by β-catenin, as detected by CUT&RUN-LoV-U (Pagella et al., 2022). Color scale indicates the relative mRNA expression level, while the dot diameter is proportional to the fraction of cells that express each gene

## Methods

### Data availability

All datasets described in this work are publicly available at ArrayExpress (https://www.ebi.ac.uk/arrayexpress/) with accession number E-MTAB-12598.

### hESCs culture and treatment for scRNA-seq

hESCs were cultured in Essential 8™ medium (A1517001, ThermoFisher Scientific) in plates and flasks coated with 10 μg/ml Vitronectin (A14700, ThermoFisher Scientific). Cells were passaged by incubation in 0.5 mM EDTA-PBS (stock: 0.5 M EDTA, AM9260G, Thermo Fisher Scientific). hESCs were seeded at a density of 10^4^ cells/cm^2^ in Essential 8™ medium, and after 24 hours, they were cultured in Essential 8™ medium supplemented with 10 μM CHIR99021 (SML1046-25MG, Sigma-Aldrich). Cells were then detached after 4 hours, 24 hours, and 72 hours of CHIR treatment (n = 6 wells per time point). The cell suspensions were filtered through a 40 μm cell strainer (KKE3.1, Carl Roth), and cell concentrations were evaluated using a hemocytometer in a Leica DM 204 IL LED microscope and adjusted to 1”000 cells/µl. 10”000 cells per sample were loaded into the 10X Chromium controller and library preparation was performed according to the manufacturer”s indications (10X Genomics; v3 protocol). Libraries were validated on an Agilent 2100 and quantified using quantitative PCR (Q-PCR). Libraries were then sequenced on Illumina NovaSeq 6000 S4 flow-cell with PE150 according to results from library quality control and expected data volume.

### ScRNA-seq data analysis

Raw sequencing data in FASTQ format were processed with 10X Genomics Cell Ranger 7.0.1 for read alignment and feature-barcode matrix generation using the cellranger count pipeline. Human reference genome GRCh38 2020-A was downloaded from the 10X Genomics homepage (https://support.10xgenomics.com/single-cell-gene-expres-sion/software/overview/welcome).

Subsequent analyses were performed in the programming language R (version 4.0.3) and Rstudio (version 1.1.463). Cell Ranger output feature-barcode matrices in HDF5 format were loaded into R and processed using the Seurat package (version 3.2.3), a comprehensive framework for single-cell omics analysis developed by the Satija Lab (Stuart et al., 2019). Cells with less than 200 detected genes were removed, together with cells with more than 20% mitochondrial reads and/or less than 5% ribosomal reads. Also, genes expressed in less than 3 cells were removed from the data.

Filtered data were normalized for sequencing depth using the Seurat NormalizeData function with LogNormalize method and default settings. Next, as scRNA-seq data commonly contain many false zeros, missing values were imputed using Adaptively-thresholded Low Rank Approximation (ALRA) (Linderman et al., 2022), bringing the data from about 27% non-zero values to about 45% non-zero values. The top 3000 highly variable genes across cells were identified with the Surat FindVariableFeature function using a variance stabilizing transformation (VST) method. These variable genes were used for subsequent data scaling and dimensionality reduction steps. In addition, doublets among the cells were predicted and removed using DoubletFinder (McGinnis et al., 2019). Analyzed datasets were integrated using the Harmony algorithm (Korsunsky et al., 2019). Differential gene expression between timepoints was assessed using the Seurat FindAllMarkers function with Wilcoxon Rank Sum test (logfc.threshold = 0.25, min.pct = 0.1, min.diff.pct = 0.2, only.pos = FALSE). Differential gene expression between selected groups of cells (*AXIN2*^*high*^ */ SP5*^*low*^ and *AXIN2*^*low*^ */ SP5*^*high*^) was assessed using the Seurat FindMarkers function with DESEq2 method (logfc.threshold = 0.25, min.pct = 0.1, min.diff.pct = 0.2).

### Human healthy intestine and colorectal cancer (CRC) scRNA-seq dataset analysis

The CRC scRNA-seq data were obtained from https://sys-bio.net/sccrc/ (Uhlitz et al., 2021) and processed through the standard analysis steps The analysis was performed using Seurat (version 3.2.3) and R version (version 4.0.3). For further details about the data analysis see previous section above and attached code.

### β-catenin CUT&RUN-LoV-U data

β-catenin time-resolved CUT&RUN-LoV-U data were obtained from ArrayExpress ((https://www.ebi.ac.uk/arrayex-press/), accession number E-MTAB-12077 (Pagella et al., 2022). The protocol for CUT&RUN-LoV-U against β-catenin is described in detail in (Zambanini et al., 2022).

